# Disentangling the age-dependent causal pathways affecting multiple paternity in house sparrows

**DOI:** 10.1101/2024.02.05.579001

**Authors:** Jørgen S. Søraker, Peter Sjolte Ranke, Jonathan Wright, Thor Harald Ringsby, Bernt-Erik Sæther, Henrik Jensen, Yimen G. Araya-Ajoy

**Affiliations:** Centre for Biodiversity Dynamics, Department of Biology, Norwegian University of Science and Technology, Høgskoleringen 5, NO-7491 Trondheim, Norway; Edward Grey Institute, Department of Biology, University of Oxford, Oxford, UK; BirdLife Norway, Sandgata 30 B, NO-7012 Trondheim, Norway

**Author notes:** Correspondance: Jørgen S. Søraker.

**Keywords:** age, badge size, extra-pair paternity, house sparrow, plasticity, secondary sexual trait, selective disappearance

## Abstract

The causes of variation in multiple paternity (MP) in socially monogamous birds have received considerable attention. Traits like age and age-dependent morphology have been shown to be important for the distribution of MP. However, most studies fail to separate between the different processes underlying age-effects on MP, such as selective disappearance (that is, higher survival probability for certain phenotypes) versus individually plastic changes in morphology and mating behaviour with age. Using Bayesian multi-level path analysis on a long-term dataset from a house sparrow (*Passer domesticus*) metapopulation, we disentangle the effects of age on male and female MP, both independently from and through morphological traits. Age was a key determinant of MP for males when accounting for morphological traits, with males having MP more often as they get older (plastic component of age) and not due to disappearance of males unsuccessful to get MP (selective disappearance), challenging ‘good genes’ explanations for MP. Conversely, selective disappearance explained the higher levels of MP in older females, suggesting that higher quality females have MP more often than lower quality females. We show how an appropriate statistical decomposition of age-components and age-dependent processes provides insights to the biological drivers of MP.

## Introduction

Understanding natural variation in mating systems has been a major undertaking in ecological research over the past decades. In particular, the study of multiple paternity (MP), where an individual female reproduces with multiple male individuals at the same time, has received considerable attention because it affects sexual selection and population dynamics (i.e. through increasing variance in reproductive success and effects on effective population size [1–3]). In birds, MP has mainly been investigated either as social polygamy, sequential polygamy or through extra-pair paternity (EPP) [4]. Where the drivers of which individuals can engage in these behaviours can be expected to be similar.

MP has been intensively studied in socially monogamous bird species to understand why females engage in such behaviour [5–7]. Trivers [8] suggested that males should seek extra-pair copulations to maximize the number of offspring, while females should seek extra-pair copulation to increase offspring genetic quality. Since then, suggested hypotheses describing the potential benefits of this behaviour for females include: (a) ‘good genes’ where offspring fitness is improved by higher quality or attractive male genes (including the ‘sexy son’ hypothesis [9]), that females identify from male phenotypic cues [4,10,11]; (b) ‘compatible genes’ where offspring fitness increases due to a better genetic match between their mother and the extra-pair male [12–14]; (c) ‘fertility assurance’ by increasing the chance of females producing fertilized eggs [15,16]; and (d) MP as a form of genetic bet-hedging under unpredictable environments [17]. However, there might also be costs associated with multiple mating, such as reduced paternal care to the brood, search costs for additional partners and parasite transmission [18].

Individuals involved in MP are not expected to be a random subset of the adult population [19], and traits correlated with MP in both males and females have attracted considerable attention [19–21]. In particular, male secondary sexual traits have been widely investigated in terms of how they affect total mating success [19,21–23]. However, their relative importance has been brought into question by the possible effects of publication bias [24,25]. Moreover, few studies have controlled for age-dependent confounds in these secondary sexual traits, which could also result in misleading conclusions.

For example, the size of the black breast plumage bib, or ‘badge’, in male house sparrows (*Passer domesticus*) has been used as an example of a status signal after studies revealed larger badges were associated with higher levels of EPP [20,26]. However, other studies have failed to find support for such a relationship [27]. Meta-analysis has since demonstrated publication bias towards positive results regarding the effects of male house sparrow badge size as a status signal [25]. Moreover, badge size in male house sparrows is age-dependent, with older age classes generally displaying larger badge sizes [28–31]. As age has been found to be an important factor in determining total mating success [32], studies may draw the wrong inferences if the effects of badge size and age are not properly decomposed.

Age has been found to be a key factor predicting both the loss of within-pair paternity by young males and the acquisition of EPP by older males [32–34]. However, the expected among-individual correlation between the loss of within-pair paternity and gaining of extra-pair paternity turns out to be non-significant across species in general [32,35,36]. For example, male house sparrows in older age classes gained more extra-pair offspring, but not more extra-pair copulations, indicating a role for (possibly female mediated) post-copulatory advantage for older males [37,38]. Older males may also be better at obtaining copulations because of experience, or they might be better at convincing or forcing copulations with females, known as the ‘manipulation hypothesis’, which thus predicts that older and larger males will obtain more MP [7,34]. Meanwhile, fewer studies have investigated the role of female age. Studies have found support for both a positive relationship between female age and MP [39], as well as a negative relationship [40]. Therefore, the role of age and the underlying age-dependent processes are even less understood for females.

Although studies have investigated the role of age, there are different underlying processes by which age could influence MP. Male age could operate through increased within-individual reproductive investment with age, or through among-individual selective disappearance, where individuals that survive to reach older age classes tend to be higher quality individuals [41]. However, not many long-term studies have quantified the relative importance of within-individual plasticity with age versus selective disappearance in wild populations [42,43]. A recent study of the cooperative breeding Seychelles warbler (*Acrocephalus sechellensis*) indicates that within-individual plastic changes in mating behaviour were the main driver behind the effect of age on EPP [43]. Conversely, Segami et al. [44] found that among-individual effects of age due to selective disappearance were most important for EPP in collared flycatchers (*Ficedula albicollis*). Nevertheless, the combined roles of these different types of age effects and (possibly age-dependent) secondary sexual traits remain poorly understood.

In addition to morphology and age, other factors could also be important drivers of rates of MP, such as the breeding synchrony of individual females (i.e. the timing of breeding relative to the other females in the population; [45]). Breeding synchrony can affect levels of MP in several ways. First, it can decrease the availability of potential extra-pair mates because males could be busy performing paternal care or mate guarding [4]. Second, it can increase the level of MP because it facilitates female comparisons of different males [45]. Either way, these population-level effects of breeding synchrony on MP have proved difficult to demonstrate [26,46].

The objectives of the current study was to quantify the relationships between age and age-dependent morphological traits, and their relative contributions in explaining individual variation in MP in males and females using long-term data from a house sparrow metapopulation. The house sparrow is mainly socially monogamous, and occasionally socially polygynous [47]. Therefore, for males, MP is defined as siring offspring in multiple nests simultaneously (within the time it takes to complete a clutch), and as having more than one male siring offspring within the same clutch for females. We investigated the underlying within-versus among-individual effects of age by explicitly modelling selective disappearance versus age-related plasticity both with respect to morphology and to MP. Using multi-level Bayesian path analysis, we quantified the causal pathways linking MP with age and morphology at the among- and within-individual level for both males and females. This multi-level parametrization of path analysis allowed us to study the independent and total effects of morphology and age on MP that were associated to selective disappearance and within-individual plasticity.

## Methods

### Data collection

We used data from a long-term study of house sparrows in an insular metapopulation on the Helgeland archipelago (66.30°–66.80°N, 12.00°–13.10°E), in northern Norway between 1993–2014 that include 18 different islands, of which 15 were used in this study (Figure 1). The population on the island of Ytre Kvarøy went extinct in 2000 after occupying the islands for several decades [48], and the population on Aldra was established by four individuals (one female and three males) in 1998 [49]. In this study system, the house sparrow breeds mostly in farms on the islands closer to the coast, and mainly in cavities on private houses and artificial nest boxes on the islands farther from the coast [50]. The breeding season in this area lasts from early May to mid-August. Throughout the breeding season, new nests were thoroughly searched for, and previously used nest sites were visited regularly. During incubation, nests were visited two or three times to estimate first egg-laying date, assuming one egg was laid each day, and using the maximum number of eggs recorded as clutch size. After hatching, if the age of nestlings was unknown then it was evaluated based upon developmental stage and descriptions of morphological development of nestlings [51]. Nestlings in this metapopulation normally stay in the nest until age of 14-18 days after hatching, and at age 7-12 days individual measurements were taken and nestlings ringed with a unique combination of one metal and three colour rings. Based on aged nestlings, hatch date was then back-calculated. Additionally, 25 μL blood was drawn from the brachial vein for use in genetic analysis.

**Figure 1:**
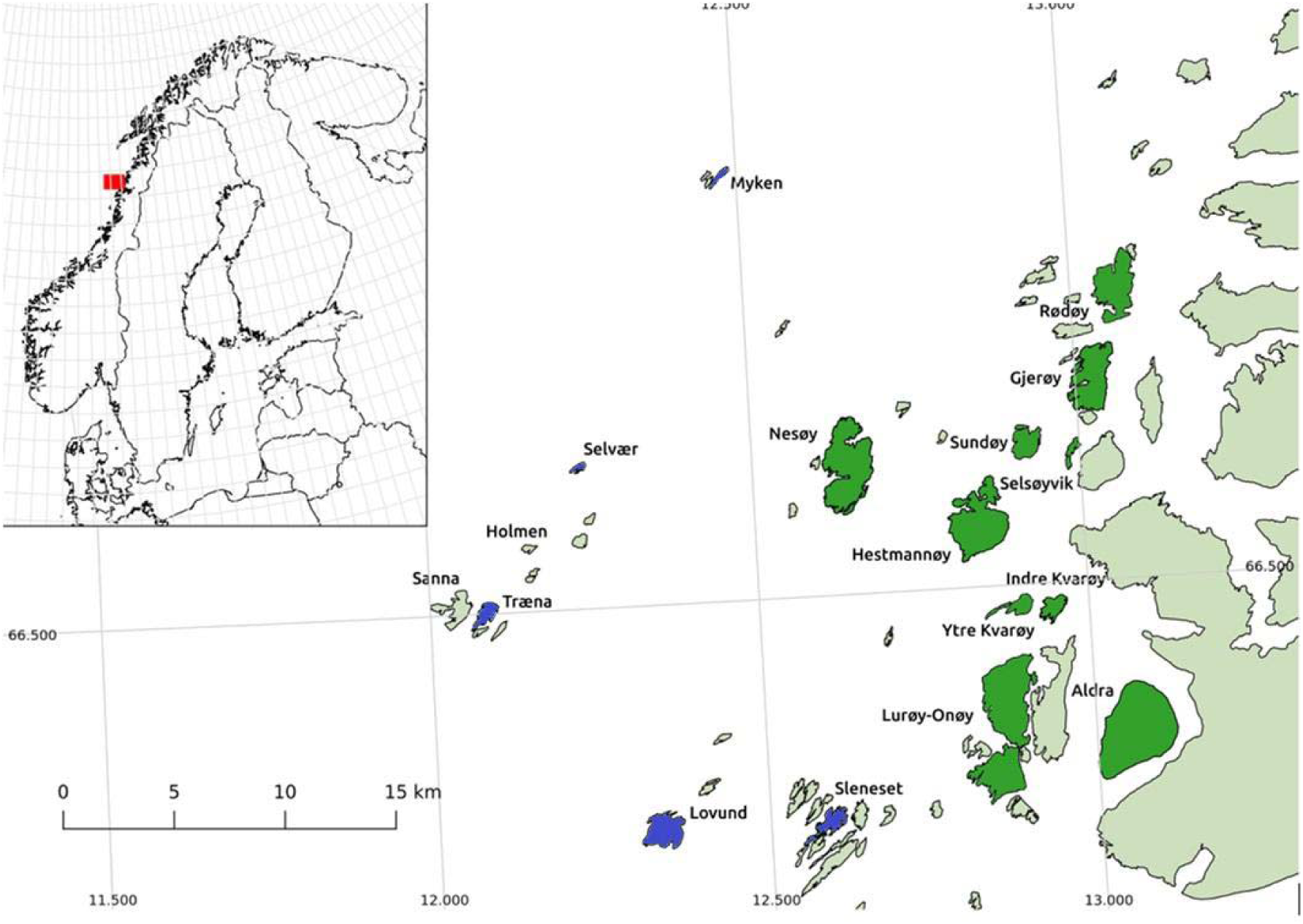
The study system in the Helgeland archipelago showing the islands included in this study. Farm islands where house sparrows usually reside close to- and on farms are marked in dark green, while non-farm islands are marked in blue. Other islands (with no house sparrows) and the mainland (with small and few populations) are marked in light green. The island populations we used MP data for in the current study were Lovund, Træna, Aldra, Selvær, Myken, Hestmannøy, Ytre Kvarøy, Indre Kvarøy, Nesøy, Gjerøy, Rødøy, Sundøy, Selsøyvik, Sleneset and Lurøy-Onøy.

Fledged juveniles and adults were caught using mist nets and morphological measurements collected. Furthermore, blood samples were collected for previously non-marked individuals, and they were ringed with a unique combination of one metal and three colour rings.

The percentage of marked adults on all islands often exceeded 80% [52,53], and the average recapture-rate on the main islands in the study system was 0.8 [54]. The average genetically determined (see below) age-at-last-reproduction (ALR) for males in this study was 1.95 years, which is likely similar to the average male lifespan. Dispersal between the islands in this metapopulation occurs regularly, with around 10.6% of the recruits on an island being dispersers [52,55]. Virtually all dispersers are juvenile [52,55], which imply that each adult can only produce offspring on one island.

### Pedigree

Genetic parentage was determined for clutches in the 15 largest insular house sparrow populations in our study metapopulation. We used the number of identified genetic fathers within a clutch to confirm the presence-absence of MP. Hence, we did not distinguish between MP for clutches with two or more genetically assigned fathers. In this metapopulation, the sparrows can have up to three clutches within a season. To assign genetic parentage of the sparrows, we extracted DNA from blood samples of ringed nestlings and adults. Next, individual genotype data generated by polymerase chain reaction (PCR) amplification of up to 14 polymorphic microsatellites was used in the parentage assignment software CERVUS [56]. Females were assigned MP if their clutch had multiple sires in at least one clutch for that year (yearly overview given in Table S1). Males, on the other hand, were assigned MP if they were assigned offspring in at least two different clutches of different females with hatch dates less than 30 days apart (i.e. a full breeding cycle; yearly overview given in Table S2). Since breeding events are not equally or normally distributed across the breeding season due to multiple clutches, the same analysis was also performed using only first clutches for females to better investigate the role of breeding synchrony. Using a subset of clutches with known social parents (n=97), we corroborated that MP equals EPP in 82% (n=17) of the cases where the social father did not sire all offspring (n=14). Only ringed individuals were sampled for blood and genotyped. Hence, the assignment of MP is likely underestimated because egg hatching failure and chick mortality can lead to MP-offspring not being sampled. Therefore, when using the term MP, we refer to observed MP measured in terms of multiple genetic sires recorded among nestlings present in a clutch at the time of ringing and blood sampling (mean age of sampled nestlings: 9.08 days, SD: 2.56 days.

### Definition of variables

#### Breeding synchrony

To investigate how synchrony (through nest hatch date relative to other nests in a specific island and year) influenced the occurrence of MP for females, the hatch dates were mean centered within each year and island. We only included the first clutch in these analyses since broods were not evenly or normally distributed across the breeding season due to multiple clutches, therefore being a poorer measure of available mates than if only first clutches are considered (see breeding synchrony hypothesis in the introduction).

#### Morphology

Body mass was measured in both females and males using a Pesola spring balance to nearest 0.1 g. Because body mass has previously been shown to be a reliable proxy for body size in this metapopulation [57], body mass was used as a measure of body size,.

When adult males were captured, both the height and width of their breast ‘badges’ were measured to nearest mm. Badge sizes were measured as described by Møller [58], using the ‘potential’ (total) area of black badge feathers rather than the visible ‘realized’ area. These two ways of measuring the badge show similar tendencies with age in this metapopulation, although the realized badge size is more sensitive to environmental conditions [29]. The calculation of badge size for males was performed based on equation 1 [59], which corresponds to the ‘total’ badge size according to Møller and Erritzøe ([60]):

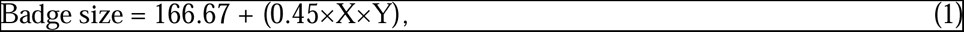

Where X is the total badge height and Y is the total badge width.

Body size has previously been hypothesized to be important for MP, since larger males could potentially attract and/or force copulations more effectively [34,61]. Moreover, body mass has been shown to be positively correlated with total badge size in house sparrows both phenotypically and genetically [30].

Morphological traits were log-transformed (natural logarithm) to explicitly model allometric relationships. Most individuals were measured across at least two years (Table S3).

### Statistical analyses

#### Pathways to MP

We used a multi-level Bayesian path analysis to study a set of *a priori* hypothesized relationships between aspects of age, morphological traits and MP. To facilitate comparisons and to provide an overview of all hypotheses, these are summarized in a path diagram (Figure 2 and 3 for males and females respectively). These relationships were then parameterized as a joint likelihood path model in a Bayesian mixed-model framework. The total effect of the predictors of MP was calculated based upon the path rules established by Wright [62] and the corresponding extensions to nonlinear relationships [63–65]. The total paths captured the overall effect associated to the direct and indirect pathways (Table S4 & S5). In this way, we aimed to describe the causal relationships between morphological traits, age, and the occurrence of MP. For example, the total effect of body mass on MP in males was calculated as the direct effect of body mass on MP, plus its indirect effect through badge size, which was calculated as the product of the effect of body mass on badge size and of badge size on MP.

**Figure 2:**
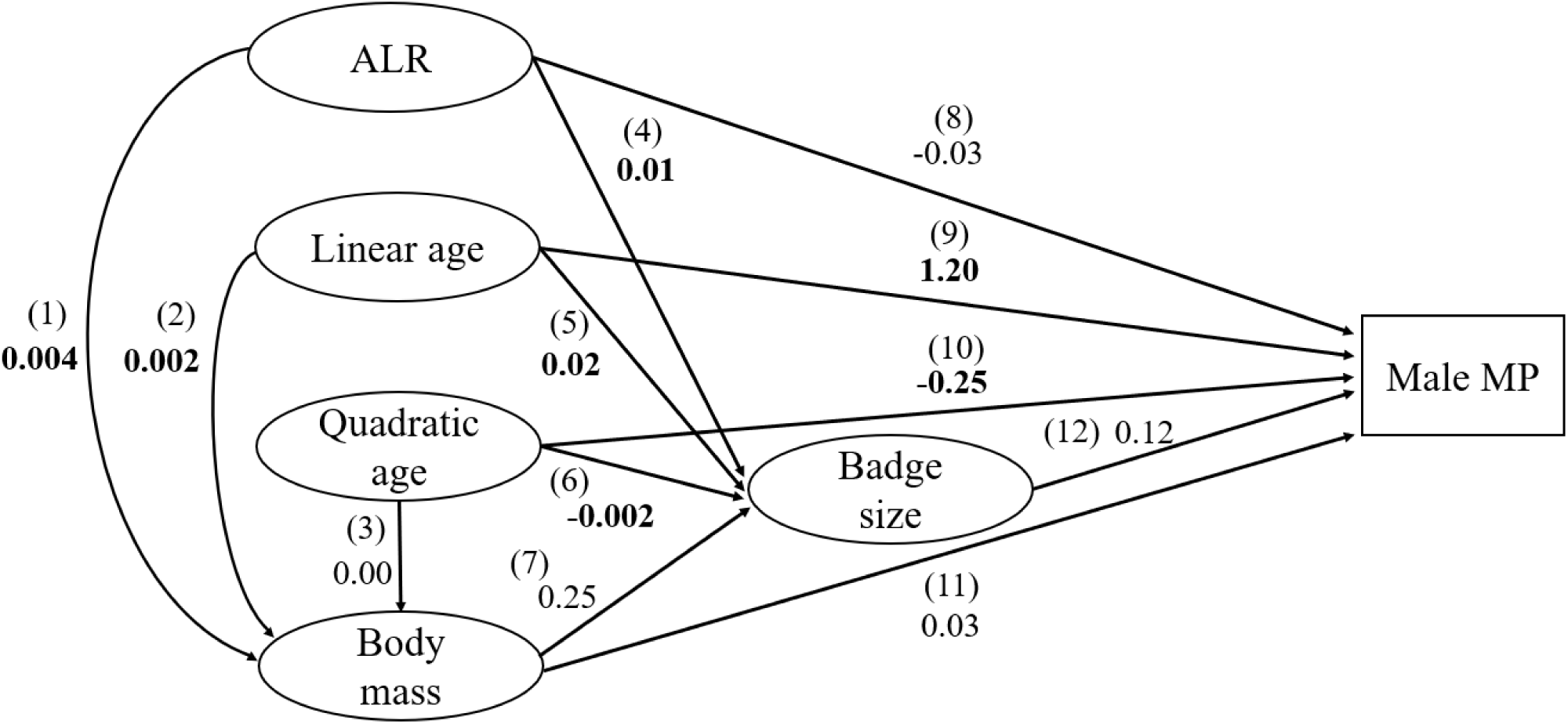
Path diagram illustrating the direct and indirect pathways of age-at-last-reproduction (ALR), age, quadratic age and age-dependent traits affecting MP for males. Path number, as described in the methods, is given in parentheses. Point estimates in bold had strong support by the model (95% CrIs did not include zero).

**Figure 3:**
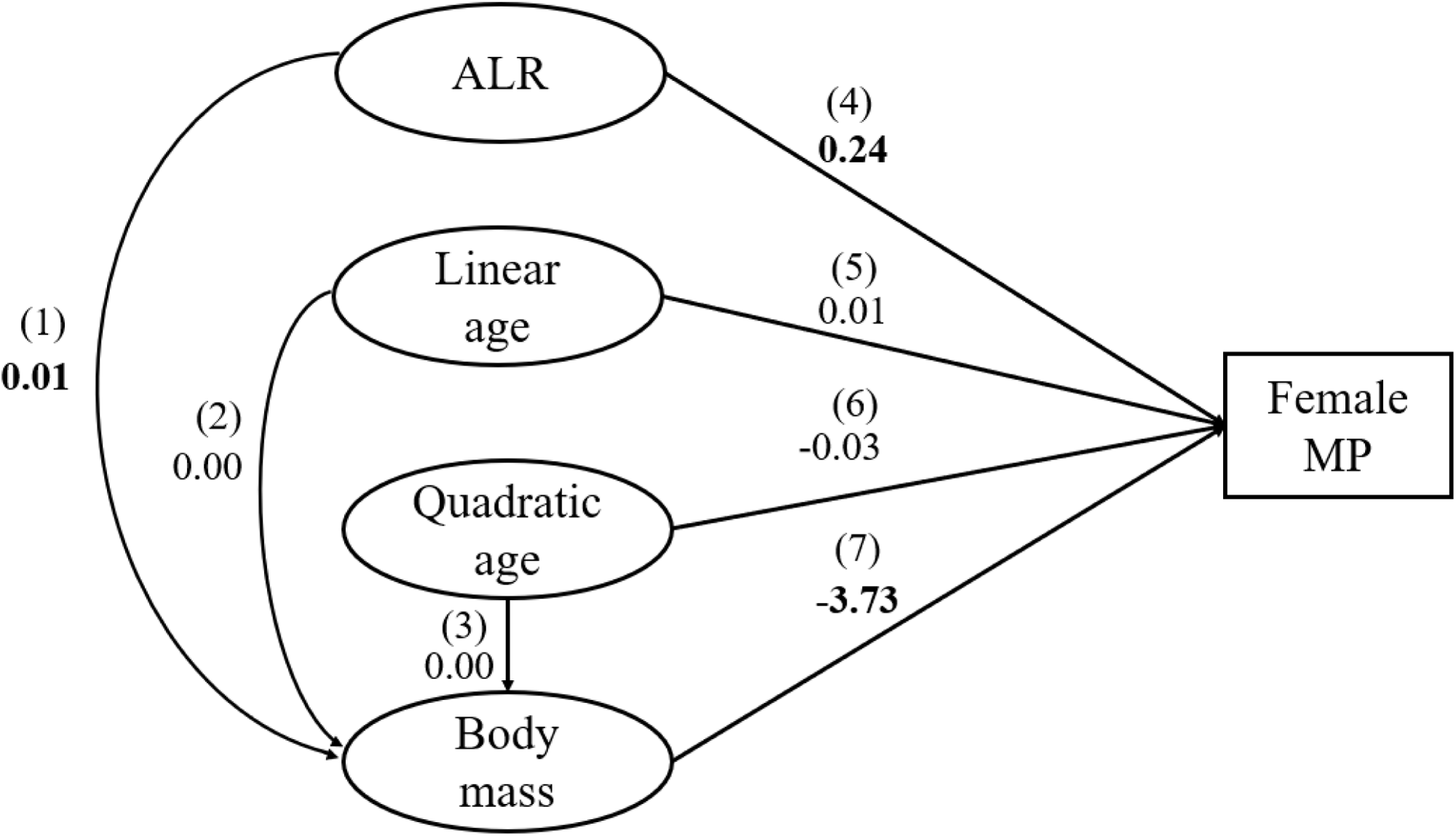
Path diagram showing the direct paths of different components of age (including age-at-last-reproduction ALR), age-dependent morphological trait and breeding synchrony and their effect on MP for females. Path number, as described in the methods, is given in

#### Statistical implementation

We used STAN through the RSTAN package [66] in the R environment [67] to estimate the joint likelihood of the different models that were part of each path analysis. This approach allowed us to obtain appropriate measures of the uncertainty of total pathways. If a morphological measurement was missing for an individual in a given year, the mean value of the individual for that morphological trait was used when the morphological trait were fitted as a response variable [68]. We were then able to correct for this missing information when the morphological measures were used as a predictor, because we used the model predictions of one sub-model (e.g., body mass sub-model) as the predictor for the subsequent sub-model (e.g., multiple paternity sub-model) instead of a missing value. To validate the robustness of this approach, we performed a simulation study based upon the results obtained in the statistical analyses and the missing data structure of our data set (see Supplementary Materials and Table S6). To validate the direct effects outside a Bayesian framework, we also corroborated the results using a stepwise frequentist approach (“lmer” package [69]). To control for the potential effects of clutch-availability, reflecting clutches the male could potentially sire offspring in, we also fitted models including the number of first clutches for each island in a given year (see Table S7 & S8).

#### Modelling age effects

Age can affect MP directly or indirectly through different paths of cause and effect. First, older individuals may be of ‘higher quality’, purely because of the selective disappearance of ‘lower quality’ males, which will cause an among-individual effect of age on MP. By ‘quality’, we here mean an individual’s survival propensity which partly reflects its expected fitness [70]. Second, individuals may improve their abilities, levels of investment and/or relative condition with age, reflected as a within-individual plastic effect of age on MP. In order to decompose the effects of age into either within-versus among-individual effects, we used age-at-last-reproduction (ALR) per individual to estimate selective disappearance [41]. We also fitted age itself to estimate the plastic linear effects associated with age, and the quadratic effect of age to test for any non-linear relationship [71]. Age and ALR was measured in years as the true age in years −1, with intercept thus representing 1-year old individuals. In this metapopulation, virtually all individuals start breeding at the age of one year, so we did not include age-at-first-reproduction to model selective appearance. Since age and ALR were on the same unit scale (age in years), the direct effects of these variables can be compared. The total paths (direct and indirect effect combined) can also be compared as their effects are still biologically relevant units. Hence, testing both for among-individual (ALR) and within-individual (age and quadratic age) changes allowed us to disentangle within-individual plasticity versus among-individual selective disappearance [41] when both were included in the different models (described in detail below).

#### Model for males

We used mixed-effect models in a path analysis framework for males based upon three different sub-models (with numbers in parenthesis here referring to path links given in Figure 2 and 3). In the first sub-model, the direct effects on log(body mass) from ALR (1), age (2) and quadratic age (3) were estimated by fitting them as fixed effects, with body mass as the response variable with a Gaussian error distribution. Estimates are interpreted by the increase in log body mass as the age predictors increase with one year. In the second sub-model, log(badge size) was treated as response variable with a Gaussian error distribution, and ALR (4), age (5), quadratic age (6) and body mass (7) were fitted as fixed effects. In the last sub-model, male MP was fitted as a response variable with a binomial error distribution, and ALR (8), age (9), quadratic age (10), body mass (log) (11) and badge size (log) (12) were fitted as fixed effects. We included a quadratic age effect to allow for a non-linear effect of age on MP, because a non-linear relationship between EPP and age has been documented in other species [32,43]. Individual identity and island-year (combination of island and year) were fitted as random effects for all sub-models. From this model, the estimates are interpreted as the change in the odd log ratio of MP as the predictor increase with one unit. The total paths were then calculated following the path rules by [62] (Table S4 & S5). This set of analyses was carried out using data from the entire breeding season and repeated using just the first clutch for each female.

Whenever both the linear and quadratic effects of age were supported by the model, we wanted to estimate age at which the trait value peaked. Since the linear effect was always positive and the quadratic effect always negative, the age with highest peak value was calculated as:

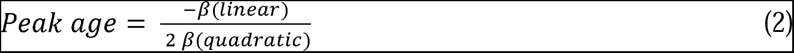

#### Model for females

The path analysis for females was based upon two sub-models using two different datasets (with numbers in parenthesis here referring to path links given in Figure 2 and 3). First, we modelled body mass as response variable with ALR (1), age (2) and quadratic age (3) as fixed effects, with Gaussian error distribution. Second, female MP was response variable with binomial error distribution, and ALR (4), age (5), quadratic age (6), body mass (7) and breeding synchrony (8) as fixed effects. This was done with the dataset including only first clutches for the females. Thereafter, the same models were fitted using all available clutches over the breeding season, while leaving breeding synchrony out of the models. All models included individual identity and island-year as random effects.

## Results

### MP in the metapopulation

In this metapopulation, 23.71% of the males (total N = 485) obtained MP at some point during their lifetime. For females (total N = 486), 43.00% had MP at least once in their lifetime. The proportion of clutches (total N = 1386) with more than one genetic father fluctuated among years (mean = 43%; range 25%-66%; N = 21 years) (Tables S1 and S2).

### Age-dependent male body mass

There was a positive among-individual relationship between ALR and body mass, indicating that heavier males were more likely to reach the older age-classes than lighter males (Table 1 and Figure 2). We also found support for a linear effect of age on body mass, implying that there was a systematic within-individual increase in adult male body mass with age.

**Table 1:** Effects of morphological traits and two age-components (including age-at-last-reproduction, ALR) on body mass, badge size and MP for males over the breeding season (Male MP). Presented estimates are the median and the 95% credible intervals (CrIs). Values in bold are supported by the model (95% CrIs not overlapping zero). The results are based on the multi-level Bayesian model composed by the sub-models described in the text.

|  | **Body mass** | **Badge size** | **MP** |
| --- | --- | --- | --- |
| **Direct effects<sup>a</sup>** | **$\beta$ [95% CrI]** | **$\beta$ [95% CrI]** | **$\beta$ [95% CrI]** |
| Intercept<sup>b</sup> | **3.45 [3.45, 3.46]** | **3.66 [3.66, 3.68]** | **-2.23 [-2.49, -1.62]** |
| ALR | **0.004 [0.003, 0.007]** | **0.01 [0.01, 0.02]** | -0.03 [-0.10, 0.21] |
| Age | **0.002 [0.001, 0.006]** | **0.02 [0.01, 0.03]** | **1.20 [0.97, 1.90]** |
| Quadratic age | 0.000 [-0.001, 0.000] | **-0.002 [-0.003, -0.001]** | **-0.25 [-0.30, -0.12]** |
| Body mass | - | 0.03 [-0.04, 0.19] | 0.03 [-0.58, 1.81] |
| Badge size | - | - | 0.12 [-0.48, 1.96] |
| **Total path** |  |  |  |
| ALR | - | **0.01 [0.01, 0.02]** | -0.02 [-0.09, 0.21] |
| Age | - | **0.02 [0.01, 0.03]** | **1.20 [0.97, 1.90]** |
| Quadratic age | - | **-0.002 [-0.003, -0.001]** | **-0.25 [-0.30, -0.12]** |
| Body mass | - | - | 0.04 [-0.60, 1.79] |
| **Random effects** | **$\sigma$ [95% CrI]** | **$\sigma$ [95% CrI]** | **$\sigma$ [95% CrI]** |
| Island-year | 0.010 [0.009, 0.014] | 0.015 [0.011, 0.028] | 0.841 [0.649, 1.337] |
| Individual | 0.056 [0.055, 0.060] | 0.098 [0.095, 0.106] | 0.650 [0.378, 1.424] |
| Residual variance | 0.031 [0.030, 0.032] | 0.070 [0.069, 0.074] |  |
| **Sample sizes** | **n** | **n** | **n** |
| Island-year | 170 | 170 | 109 |
| Individuals | 483 | 483 | 238 |
| Observations | 1024 | 1024 | 348 |

<sup>1a</sup> Estimates rounded to two decimals, unless the third decimal is needed to identify positive or negative values, in which three decimals are given
<sup>b</sup> Estimates with age 1 as reference (1-year olds set to 0) for all traits

### Age- and mass-dependent badge size for males

For badge size, we found support for the effect of age and quadratic age (Table 1, Figure S1). Furthermore, there was also a positive effect of ALR on badge size (Table 1). Therefore, there was a within-individual plastic non-linear increase of badge size with age, until a peak badge size at the age of 5 years. Although a quadratic effect was detected, it does not necessarily demonstrate a decrease in badge size after 5 years. Unfortunately, not enough data were available from older males to test for such a decrease. However, it was also the case that fewer smaller-badged individuals reached these older age classes. This was captured by the among-individual effect of ALR, demonstrating the disproportionate contribution to larger mean badge sizes of individuals that managed to survive to old age. The point estimate for body mass affecting badge size was positive, but the 95% CrI overlapped zero and we thus found no support for the expected allometric relationship between body size and badge size, after correcting for the effects of age.

Our path model also allowed us to estimate the total effect of age and quadratic age on badge size, again with badge size increasing most during younger ages as a within-individual plastic effect of age. In addition, the total path of ALR revealed its overall positive effects on badge size (Table 1). Accordingly, both among- and within-individual effects of age were found to directly affect badge size after controlling for body mass. Running the same models using a stepwise approach in a frequentist framework revealed similar results (Table S9), as did models controlling for the number of available first clutches within the given island in the given year (Table S7).

### Age and morphological traits affecting MP for males

ALR had no direct effect on MP for males (Table 1 and Figure 2). However, the model showed that age positively affected MP directly, with a negative effect of quadratic age (i.e., MP increased at younger ages, and declined at older ages). This non-linear relationship was also supported by the total path calculations. Hence, the probability of obtaining MP for males increased non-linearly with age as a result of age-specific within-individual plasticity, until peaking at an age of 3.18 (3-4) years old (Figure S2). In contrast, body mass did not have any direct or indirect effects (e.g. via badge size) on MP. Badge size showed no effect on MP (Table 1 and Figure 2). The total path of ALR and body mass was also not supported by the model. The frequentist-based model for MP, and models including the number of first clutches in the island population to correct for variation in the available nests in a given year in each island, showed similar results (Table S7 and S9).

### Age-dependent morphological traits for females

ALR positively affected body size for females (Table S10 and Figure 3), but a plastic effect of age on body mass was not supported by the model. Hence, the increase in female body mass with age was related to an among-individual effect of heavier individuals having a greater chance to reach older age. Models using a frequentist approach and controlling for number of first clutches showed similar relationships (Table S8 and S11).

### Age and morphological traits affecting MP for females

ALR was positively associated with female MP, illustrating selective disappearance of females less prone to MP. No direct effects of body mass, age or quadratic age on female MP were supported by the model (Table 2, Figure 3). Although the total path of ALR on MP included a negative estimate of body mass on MP, the effect of ALR on MP was still positive. Therefore, there was no sign of a within-individual plastic effect of age on the probability of MP for females. When modelling just the first clutches, there was no evidence for an effect of breeding synchrony within the first clutch on female MP (Table S10), we therefore found no support for the synchrony hypothesis. Frequentist models showed similar results (Table S11 and S8).

**Table 2:** Direct path coefficients of age (including age-at-last-reproduction ALR) and morphology on MP for females, using a dataset where we are not certain of the assignment of fathers. Presented estimates are the median and the 95% credible intervals (CrIs). Values in bold are supported by the model (95% CrI not overlapping zero). The results are based on the Bayesian model with sub-models described in the text.

|  | **Body mass** | **MP** |
| --- | --- | --- |
| **Direct effect**<sup>a</sup> | **$\beta$ [95% CrI]** | **$\beta$ [95% CrI]** |
| Intercept<sup>b</sup> | **3.45 [3.44, 3.45]** | 4.46 [-8.51, 17.83] |
| ALR | **0.01 [0.01, 0.02]** | **0.21 [0.07, 0.36]** |
| Age | 0.00 [-0.01, 0.01] | 0.09 [-0.33, 0.52] |
| Quadratic age | 0.001 [-0.001, 0.002] | -0.06 [-0.16, 0.03] |
| Body mass | - | -1.52 [-5.36, 2.28] |
| **Total path** |  |  |
| ALR | - | **0.19 [0.05, 0.34]** |
| Age | - | 0.10 [-0.33, 0.53] |
| Quadratic age | - | -0.06 [-0.16, 0.03] |
| **Random effects** | **$\sigma$ [95% CrI]** | **$\sigma$ [95% CrI]** |
| Island-Year | 0.02 [0.01, 0.02] | 0.26 [0.13, 0.67] |
| Individual | 0.06 [0.06, 0.07] | 0.31 [0.02, 0.76] |
|  | 0.05 [0.05, 0.05] |  |
| **Sample sizes** | **n** | **n** |
| Island-year | 186 | 119 |
| Individual | 609 | 282 |
| Observations | 1123 | 456 |

<sup>2a</sup> Estimates rounded to two decimals, unless the third decimal is needed to identify positive or negative values, in which three decimals are given
<sup>b</sup> Estimates with age 1 as reference (1-year olds set to 0) for all traits

## Discussion

Our study shows that age was an important factor in determining MP for males, in line with findings across various bird species [36]. However, most studies do not decompose the effects of the two age-related processes of within-individual age-based plasticity versus among-individual selective disappearance with age to MP [43]. In our study system, older males obtained paternity in two or more clutches simultaneously through a plastic within-individual age-effect, independent of any among-individual effects of ALR and morphology. Interestingly, badge size did not appear to influence MP, contrary to expectations from hypotheses that badge size acts as a quality and/or dominance correlated trait [20]. In addition, there was no effect of ALR through selective disappearance affecting male MP. Hence, we show that disentangling the statistical effects of male age into different ecological age-related processes can provide valuable information regarding the relative importance of these types of mechanisms in wild populations.

The finding that a plastic effect associated to age is an important driver for males in obtaining MP, is informative regarding why females might seek MP via EPP. If females seek EPP for ‘good genes’ (Westneat, 1990), one might expect high-quality males to consistently obtain more EPP. However, we found no support for ALR, an among-individual effect of age on MP, thus showing no evidence for an effect of male quality through selective disappearance on MP. This is in accordance with for example what Raj Pant et al. [43] found in Seychelles warblers (*Acrocephalus sechellensis*). Contrary to the expectation from the ‘good genes’ hypothesis, we found the within-individual plastic effect of age to be an important factor, illustrated by the evidence for a positive, but non-linear plastic effect of age. This alone does not exclude the hypothesis regarding genetic benefits, as any compatibility effect may have been obscured due to the stronger effect of a plastic component of male age. Moreover, it is possible that females choose ‘high-quality’ males using age as a proxy. A within-individual plastic effect of age, rather than selective disappearance, may reflect a different underlying process driving MP. For example, increased male investment in searching for and/or harassing females for MP, or a trade-off between within-pair paternity and extra-pair paternity [35]. Unfortunately, we cannot separate between these two hypotheses in this study. However, male house sparrows have also been found to increase within-pair paternity with age [32], indicating that increased male investment in both types of paternity may be the most feasible explanation. It should be noted that the support for a quadratic effect of age also may suggest reproductive senescence if MP is positively correlated with reproductive output [32,71]. However, due to a lack of reproductive events after the calculated peak, a post-peak decline in MP could not be formally tested for. However, a non-linear quadratic effect of age on reproduction (measured as recruits produced per year) has previously been demonstrated in this metapopulation [54]. Although previously assumed that senescence could rarely be observed in the wild, more recent evidence suggest that it is widespread and often detectable [73,74]. Our results may indicate that reproductive senescence could partly act through multiple paternity.

As MP was not related to body mass for males, this partly challenges the ‘manipulation’ hypothesis, where larger males are more able to force copulations with females [34,61]. Nevertheless, the ‘manipulation’ hypothesis includes the possibility that older males may instead have improved time-management during breeding periods, for example through experience or increased frequency of copulations, regardless of any effect of body mass, and this might constitute the underlying mechanism determining house sparrow EPP rates [34,61]. It remains unknown if the plastic component of age reflects increased male mating investment, reproductive trade-offs or simply the process of improved optimal time allocation. Meta-analyses that included studies on multiple species of birds have indicated that larger males often obtained more EPP than smaller males [24,34]. However, although both body size and badge size have previously been demonstrated to affect mating success in this metapopulation [30], this was in an analysis that did not correct for the effect of age. Akçay & Roughgarden [24] and Hsu et al. [34] also concluded that the genetic benefits alone fail to explain the distribution of EPP in birds. This is consistent with our findings here, particularly as the effects of both ALR and badge size were not supported by the model explaining MP, which would have been expected if there was an effect of ‘good genes’ on MP and if badge size was an honest signal of male ‘quality’. This does not completely rule out genetic benefits of MP, including the ‘good genes’ hypothesis, as cues regarding male quality could work through other (unmeasured) traits and does not necessarily act through increased survival probability of individuals alone. Hence, our results suggest that the ‘manipulation’ hypothesis could be a driver for coerced EPP in this study system. However, this is not necessarily evidence of coercion, as females showed constant MP while males simply obtained more MP with age. In addition, we know from other species that females can regulate and choose EPP [75], so it is unlikely that MP in this system is completely male-driven [76]. Therefore, the current study cannot be interpreted as evidence for the manipulation hypothesis alone.

When controlling for among- and within-individual effects of age, badge size did not appear to increase the chances of a male obtaining MP (Table 1), although the point estimate was positive with a skewed posterior distribution toward positive values. Badge size has been shown to be important for lifetime reproductive success [28], as well as mating success and recruit production [30] in this metapopulation (although age was included in these analyses, the underlying processes of any age effects were not examined). Hence, the effect of badge size on lifetime reproductive success previously demonstrated in this metapopulation may not be due to more frequent MP. Although badge size has been used as example of an important secondary-sexual trait [20], recent meta-analysis studies have shown no effect of badge size as a status symbol [25] or an effect of badge size on MP [32] in house sparrows. ALR was positively associated with badge size in this study, indicating that high quality individuals have larger badge sizes. This suggests that badge size may be a proxy for male quality in this metapopulation. Although leaving out any age component in the analysis did not change the interpretation of the effect of badge size or body mass on MP (Table S12), failing to correct for age-dependency can potentially lead to misleading conclusions in other datasets [41].

The results of this study show a positive relationship between the two age parameters (age and ALR) and body mass for males. This indicates that two processes are acting simultaneously. First, males get heavier/larger as they get older through a linear effect of age. Second, the selective disappearance of lighter/smaller males contributes to higher body mass in individuals in older age classes, in line with previous results showing positive survival-selection on body mass in this metapopulation [30]. Badge size increased both with ALR and non-linearly with age. Therefore, both selective disappearance and within-individual plasticity led to larger badge sizes in older males. This demonstrates that among- and within-individual effects of age can act simultaneously. Moreover, it opens the possibility that females may use badge size as a cue for male age, although the effect of age on badge size was rather small. The effect of body mass on badge size was positive, but the credible intervals overlapped zero. This is interesting, given that body mass has been found to be a reliable proxy for body size in this metapopulation [57]. In this study, we found no evidence for a direct link between body size and badge size, two traits that have previously been shown to be positively correlated both phenotypically and genetically in this metapopulation [30]. The absence of body mass as an important factor in determining MP for males is particularly interesting. The manipulation hypothesis could predict that larger males should be better at forcing copulations or convincing females to copulate. However, we found no such support of an effect of body mass/size. This indicates that males gaining more experience in obtaining additional mating with age is the most probable explanation for why males gain more MP as they get older [34].

For females, ALR affected body mass, demonstrating that heavier/larger females were more likely to reach older age classes than lighter/smaller females via the process of selective disappearance. Neither age nor quadratic age affected female body mass, suggesting no systematic increase in female size with age. Although females have more variable body mass due to stages of egg-production, this is unlikely to drive this absence of a plastic age effect, as most measurements happened outside this period. Older females had MP more often than younger females, through the effect of selective disappearance. Hence, the occurrence of relatively more frequent MP in older females is because some ‘high quality’ females that often engage in MP were more likely to reach those older ages. Although this suggests that males might actively choose to copulate with higher quality females, it could also be that high quality females are better at turning extra-pair copulations into MP. Neither body mass, age, quadratic-age or breeding synchrony of first clutches were shown to affect MP. The lack of an effect of body mass is perhaps unsurprising, as we only investigated the role of body mass between individuals. However, within-individual relative body mass has previously been found to affect MP in pied flycatchers (*Ficedula hypoleuca*), likely through the time needed to forage versus obtain EPP [77]. The absence of any effect of breeding synchrony on female MP in the first clutch per year in this house sparrow metapopulation (measured per island per year) is interesting given that it has been found to be a driver in other bird species via the availability of extra-pair mating opportunities [45]. However, our results are in concordance with other studies that found no effect of breeding synchrony on MP (e.g. in great reed warblers (*Acrocephalus arundinaceus*) [78]). If a female breeds close in time to the breeding onset of other females, this could enable easier comparison of their social male against other alternative males, and therefore potentially increase the level of MP [45]. Simultaneously, it might enhance mate guarding from pair males and lower the level of MP [79]. It is always possible that these two processes act simultaneously to exactly balance each other, resulting in no observed effect of breeding synchrony, but breeding synchrony could also not be important for female MP in this system. This hypothesis is perhaps better explored in studies with additional data on both mate availability and mate guarding behaviour.

From an evolutionary ecology point of view, the variation in MP, particularly for males, is also interesting given that variance in reproductive success has been shown to affect effective population size in this metapopulation and other house sparrow populations along the Norwegian coast [80,81]. MP therefore appears to be an important population-level factor contributing to the rate of loss of genetic variation due to genetic drift and inbreeding, and the relative importance of genetic drift and selection for evolutionary change [82]. Thus, MP may affect temporal changes in both adaptive and non-adaptive genetic variation within and across island populations in our study metapopulation.

## Conclusions

Our study shows that older males obtained more multiple paternity (MP) because of a within-individual increase in MP with age. No evidence was found for selective disappearance as the driver behind this relationship between male age and MP. In addition, age but not body size was positively related to male badge size, through both within-individual age-based plasticity and among-individual selective disappearance. However, badge size was not associated with MP, whether or not the models controlled for the effects of male age, yet again bringing into question the signal of status hypothesis regarding badge size and MP in house sparrows. Females had more MP in higher age classes due to a selective disappearance effect of age, while no effect of an increase in age, body mass or breeding synchrony being supported to affect MP. This study emphasises the utility of statistically disentangling the different biological effects of age and morphology through path analysis to properly investigate the causal pathways of female choice and male-male competition across different types of mating systems. Thus, understanding the mechanisms for variation in reproductive success caused by MP may be important to inform us on eco-evolutionary processes.

## Authors contributions

JSØ and HJE conceived the idea. JSØ, YGA and JW designed the methodology. THR, HJE, PSR, YGA and JSØ collected the data. HJE generated part of the pedigree. JSØ and YGA analysed the data. JSØ led the writing of the manuscript, with input from all the authors. All authors contributed critically to the drafts and approved on the final manuscript.

## Acknowledgements

We want to thank Jane Reid and David Westneat for helpful comments on this study. We also want to thank all the fieldworkers that have participated in the sparrow project fieldwork, laboratory technicians who helped with genotyping for parentage analyses, and the inhabitants on Helgeland for their hospitality. This study was supported by the Research Council of Norway (RCN, project numbers 302619and 325826) and through RCN’s Centres of Excellence funding scheme (project number 223257 to the Centre for Biodiversity Dynamics (CBD) at NTNU).

