## Supplementary materials for "Disentangling the age-dependent causal pathways affecting multiple paternity in house sparrows"

**to**

**sparrows**

**Figures**

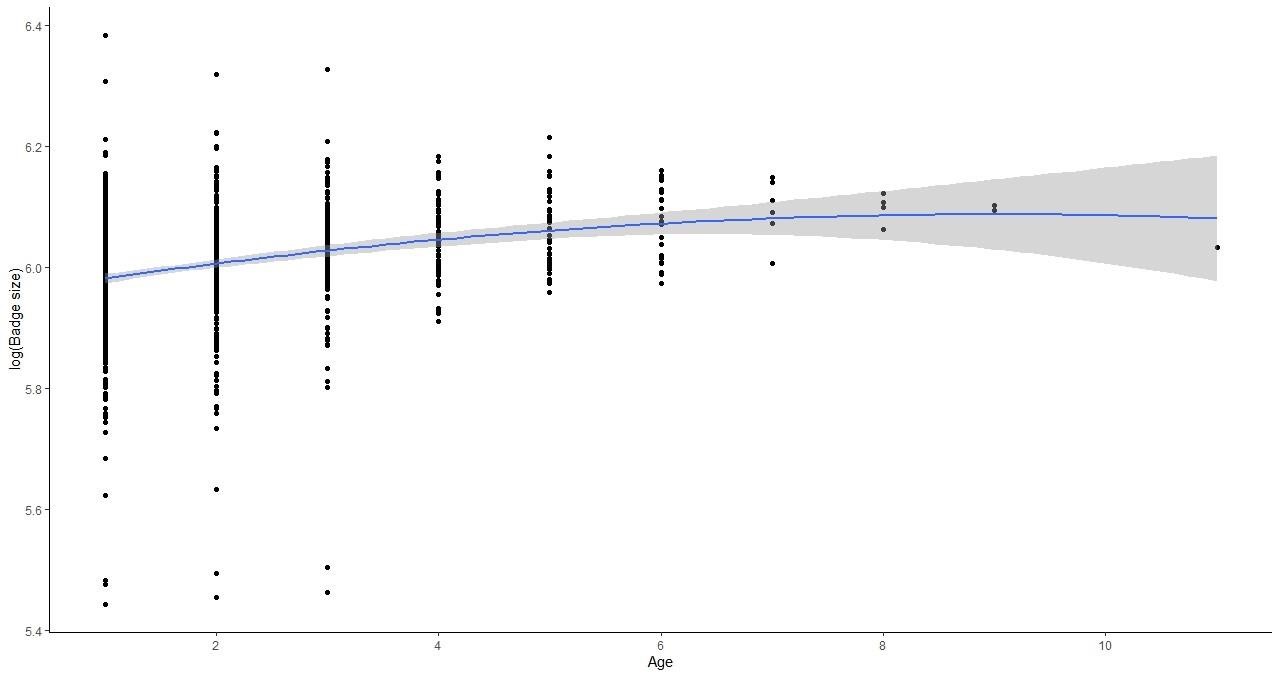

**Figure S1:** Effect of age on log-transformed badge size for males based upon the frequentist model. Here, each individual has a marginalized predicted badge size for a given age and does not take random effects of individual and island-year into account, causing the deviation from the calculated peak based on the Bayesian approach. Shaded area represents 95 % CI.

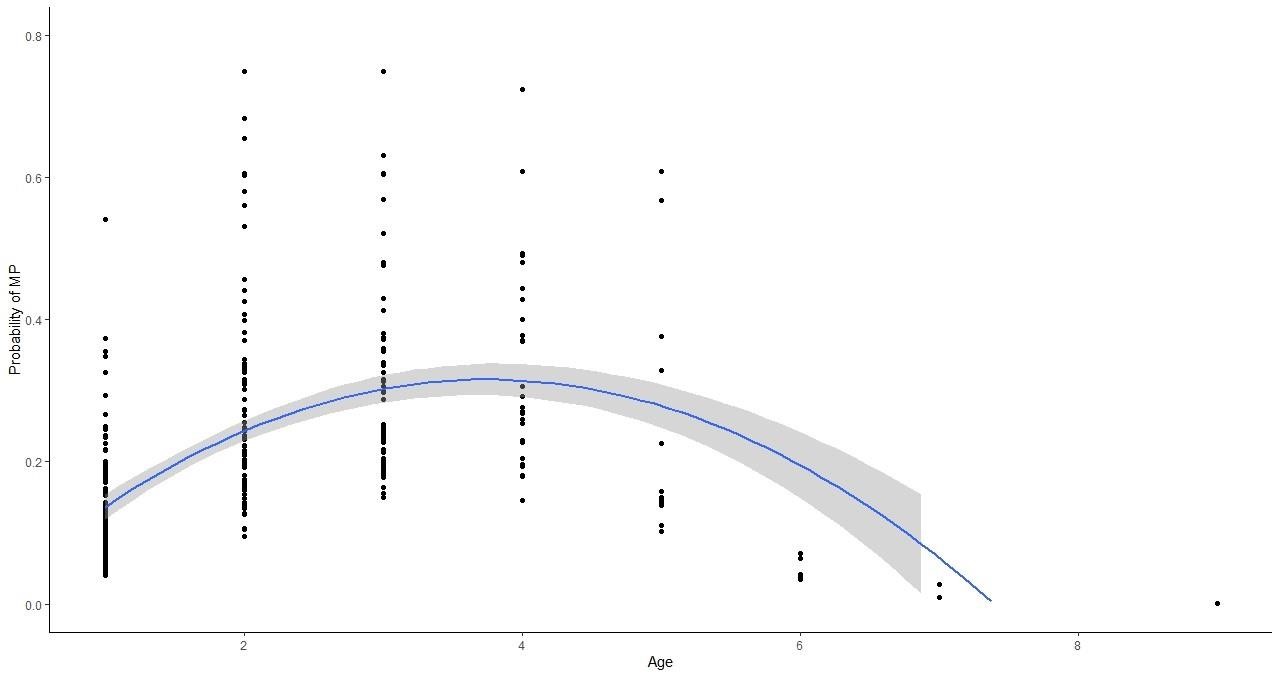

**Figure S2:** Effect of age on the probability of obtaining multiple paternity in the females first clutches (MP) for males, predicted from sub-model 3 with a frequentist approach. Each male here has a marginalized predicted value of probability of MP, which does not take the random effects of individual and island-year into account. Shaded area represents 95% CI.

**Tables**

In addition to a Bayesian framework, we also used a frequentist approach for the direct path coefficient based on the same models as for the Bayesian framework, using the “lme4” package (Bates et al. 2007). For models with mass and badge size as response-variables, we used the “lmer”-function with Gaussian error distribution, as they are both continuous variables with approximately normally distributed residuals. For the models with multiple paternity as response-variables we used the “glmer” function with binomial error distribution

**Table S1:** Yearly count of number of individually identified reproducing females, and the number of females with multiple paternity (MP) in their nest, irrespective of whether morphological data were available or not. *Number of females with MP*

| Year | Number of females with MP | Number of reproducing females |
| --- | --- | --- |
| 1993 | 17 | 41 |
| 1994 | 25 | 56 |
| 1995 | 22 | 45 |
| 1996 | 19 | 37 |
| 1997 | 17 | 41 |
| 1998 | 17 | 40 |
| 1999 | 22 | 52 |
| 2000 | 16 | 43 |
| 2001 | 24 | 45 |
| 2002 | 32 | 48 |
| 2003 | 11 | 36 |
| 2004 | 23 | 61 |
| 2005 | 19 | 76 |
| 2006 | 24 | 72 |
| 2007 | 31 | 75 |
| 2008 | 29 | 76 |
| 2009 | 40 | 86 |
| 2010 | 34 | 108 |
| 2011 | 47 | 125 |
| 2012 | 37 | 104 |
| 2013 | 31 | 91 |
| 2014 | 14 | 28 |

**Table S2:** Number of males with genetically sired offspring and number of males with multiple paternity (MP) over the study period, irrespective of whether morphological data were available or not.

| Year | Males with MP | Males with sired offspring |
| --- | --- | --- |
| 1993 | 11 | 51 |
| 1994 | 13 | 60 |
| 1995 | 10 | 50 |
| 1996 | 7 | 47 |
| 1997 | 7 | 40 |
| 1998 | 13 | 51 |
| 1999 | 12 | 57 |
| 2000 | 9 | 50 |
| 2001 | 9 | 55 |
| 2002 | 25 | 63 |
| 2003 | 10 | 53 |
| 2004 | 16 | 64 |
| 2005 | 9 | 83 |
| 2006 | 20 | 81 |
| 2007 | 21 | 99 |
| 2008 | 16 | 93 |
| 2009 | 22 | 95 |
| 2010 | 28 | 125 |
| 2011 | 40 | 155 |
| 2012 | 27 | 109 |
| 2013 | 16 | 101 |
| 2014 | 10 | 42 |

**Table S3**: Overview over how many years each individual (both males and females) was measured (repetitions). Within a year, the individual can be measured several times, in which the mean of the measurements was used.

| Repetitions | 1 | 2 | 3 | 4 | 5 | 6 | 7 | 9 |
| --- | --- | --- | --- | --- | --- | --- | --- | --- |
| Number of individuals | 284 | 172 | 82 | 37 | 26 | 9 | 1 | 1 |

**Table S4:** Direct and compound paths calculation based upon the different sub-models (on body mass, badge size and multiple paternity for Male multiple paternity, MP). The values correspond to the parameter estimates described in the text and further to the links numbered in Figure 2. The compound paths are calculated as the direct effect plus the product of the parameter estimates in the other paths.

| ***Direct effects*** | ***Body mass*** | ***Badge size*** | ***Male MP*** |
| --- | --- | --- | --- |
| ALR | 1 | 4 | 8 |
| Age | 2 | 5 | 9 |
| Quadratic age | 3 | 6 | 10 |
| Body mass | - | 7 | 11 |
| Badge size | - | - | 12 |
| **Compound path** |  |  |  |
| ALR | - | 4 + 1*7 | 8 + 1*11 + 4*12 + 1*7*12 |
| Age | - | 5 + 2*7 | 9 + 2*11 + 5*12 + 2*7*12 |
| Quadratic age | - | 6 + 3*7 | 10 +3*11 + 6*12 + 3*7*12 |
| Body mass | - | - | 11 + 7*12 |

**Table S5:** Direct and compound path calculation for models of body mass and multiple paternity for females (Female multiple paternity, MP). The values correspond to the parameter estimates described in the text and further to the links numbered in Figure 3. The compound paths are calculated as the direct effect plus the product of the parameter estimates in the other paths.

| **Direct effects** | **Body mass** | **MP** |
| --- | --- | --- |
| ALR | 1 | 4 |
| Age | 2 | 5 |
| Quadratic age | 3 | 6 |
| Body mass | - | 7 |
| Breeding synchrony | - | 8 |
| **Compound path** |  |  |
| ALR | - | 4 + 1*7 |
| Age | - | 5 + 2*7 |
| Quadratic age | - | 6 + 3*7 |

*Simulation Study*

We performed simulations to address the robustness of the multilevel path analyzes approach used in the manuscript. We specifically wanted to address whether using the predicted values for the mass and badge size could bias the results. We therefore simulated 100 data sets where the effects of the different variables were the same as the effects, we estimated in the real data set. These datasets had the same structure as real data sets obtained for the Helgeland sparrow meta-population, but we had information for all the variables for all the observations. We then proceeded to remove values in a way that the resulting data sets had the same structure of missing values as the “real data set”. We then followed the same statistical procedure as the one outlined in the main text.

We showed that the estimated values are unbiased and that the sampling variance in the simulation was similar to the estimated uncertainty for the model applied to the real data (Table S5 versus Table 1)

**Table S6:** Median and confidence interval for the results of the point estimates of the statistical models applied to the simulated data sets. (Similar table to Table 1 but with the results of the simulation.)

|  | **Body mass** | **Badge size** | **MP** |
| --- | --- | --- | --- |
| **Direct effects ^a^** | ***β* [95% CrI]** | ***β* [95% CrI]** | ***β* [95% CrI]** |
| Intercept ^b^ | **3.44 [3.44, 3.46]** | **3.66 [3.65, 3.67]** | **-2.57 [-3.38, -1.87]** |
| ALR | **0.005 [0.001,0.008]** | **0.01 [0.01, 0.02]** | 0.02 [-0.36, 0.32] |
| Age | **0.001 [0.000, 0.002]** | **0.01 [0.01, 0.01]** | **1.28 [0.61, 2.04]** |
| Quadratic age | 0.000 [0.000, 0.000] | **-0.001 [-0.001, -0.001]** | **-0.26 [-0.49, -0.10]** |
| Body mass | - | 0.001 [-0.018, 0.029] | 0.09[-1.19, 1.30] |
| Badge size | - | - | 0.06 [-1.42, 1.41] |
| **Compound path** |  |  |  |
| ALR | - | **0.01 [0.01, 0.02]** | 0.02 [-0.35, 0.32] |
| Age | - | **0.01 [0.01, 0.01]** | **1.28 [0.61, 2.05]** |
| Quadratic age | - | **-0.001 [-0.001,**  **-0.001]** | **-0.27 [-0.49, -0.10]** |
| Body mass | - | - | 0.07 [-1.19, 1.30] |
| **Random effects** | **σ [95% CrI]** | **σ [95% CrI]** | **σ [95% CrI]** |
| Island-year | 0.005 [0.005, 0.007] | 0.008 [0.007,  0.010] | 0.34 [0.13, 0.80] |
| Individual | 0.056 [0.053, 0.060] | 0.098 [0.091,  0.105] | 0.42 [0.13, 0.85] |
| Residual variance | 0.004 [0.008, 0.010] | 0.009 [0.008, 0.010] |  |
| **Sample sizes** | **n** | **n** | **n** |
| Island-year | 171 | 171 | 106 |
| Individuals | 484 | 484 | 235 |
| Observations | 1025 | 1025 | 345^1^ |

^a^ Estimates rounded to two decimals, unless the third decimal is needed to identify positive or negative values,

in which three decimals are given

^b^ Estimates with age 1 as reference (1-year olds set to 0) for all traits

**Table S7:** Effects on multiple paternity in males (Male MP) and morphology by age components (including age-at-last-reproduction ALR), morphology and total number of first clutches on the given island and year. The estimates are based on a frequentist approach, using the “lme4”-package. Values in bold are supported by the model (95% CI not overlapping zero).

|  | **Body mass** | **Badge size** | **MP** |
| --- | --- | --- | --- |
| **Direct effects** | ***β* [95% CI]** | ***β* [95% CI]** | ***β* [95% CI]** |
| Intercept | **3.45 [3.43, 3.46]** | **5.78 [5.37, 6.19]** | 6.53 [-13.44,  25.77] |
| ALR | **0.005 [0.001, 0.008]** | **0.015 [0.008, 0.022]** | 0.09 [-0.12, 0.31] |
| Age | 0.003 [-0.001, 0.006] | **0.013 [0.006, 0.020]** | **0.72 [0.09, 1.26]** |
| Quadratic age | -0.001 [-0.001, 0.000] | **-0.001 [-0.003, -**  **0.0001]** | **-0.16 [-0.29, -** **0.02]** |
| Body mass | NA | 0.05 [-0.06, 0.17] | -3.09 [-7.44,  1.21] |
| Badge size | NA | NA | 0.31 [-1.95,  2.72] |
| Number of first clutches | NA | NA | 0.01 [-0.02,  0.04] |
| **Random effects** | **σ [95% CI]** | **σ [95% CI]** | **σ** |
| Island-year | 0.008 [0.000, 0.012] | 0.000 [0.000, 0.016] | 0.739 |
| Individual | 0.057 [0.052, 0.061] | 0.011 [0.010, 0.012] | 0.413 |
| Residual variance | 0.029 [0.027, 0.031] | 0.058 [0.055, 0.062] |  |
| **Sample sizes** | **n** | **n** | **n** |
| Island-year | 171 | 171 | 112 |
| Individuals | 484 | 484 | 283 |
| Observations | 1025 | 1025 | 448 |

**Table S8:** Effects on female multiple paternity MP and morphology by age components (including age-at-last-reproduction ALR), morphology, breeding synchrony and total number of first clutches on the given island and year. The estimates are based on a frequentist approach, using the “lme4”-package. Values in bold are supported by the model (95% CI not overlapping zero).

|  | **Body mass** | **MP** |
| --- | --- | --- |
| **Direct effect** | ***β* [95% CI]** | ***β* [95% CI]** |
| Intercept | 3.45 [3.44, 3.45] | 9.81 [-3.12, 23.24] |
| ALR | **0.01 [0.01, 0.02]** | **0.24 [0.01, 0.45]** |
| Age | -0.001 [-0.007, 0.005] | 0.18 [-0.45, 0.84] |
| Quadratic age | 0.000[-0.001, 0.002] | -0.09 [-0.23, 0.05] |
| Body mass | NA | -3.24 [-7.15, 0.51] |
| Breeding synchrony | NA | 0.13 [-0.19, 0.46] |
| Number of first clutches | NA | -0.003 [-0.037, 0.030] |
| **Random effects** | **σ [95% CI]** | **σ [95% CI]** |
| Island-year | 0.017 [0.011, 0.022] | 0.967 |
| Individual | 0.062 [0.057, 0.068] | 0.021 |
| Residual variance | 0.048 [0.045, 0.051] |  |
| **Sample sizes** | **n** | **n** |
| Island-year | 184 | 99 |
| Individual | 608 | 211 |
| Observations | 1121 | 284 |

**Table S9:** Direct path coefficients of age (including age-at-last-reproduction ALR) and morphology on body size, badge size and multiple paternity for males (Male MP) over the entire breeding season using a frequentist approach with the “lme4” package. Random effects of the binomial male MP model are lacking CIs. Values in bold are supported by the model (95% CI not overlapping zero).

|  | **Body mass** | **Badge size** | **MP** |
| --- | --- | --- | --- |
| **Direct effects** | ***β* [95% CI]** | ***β* [95% CI]** | ***β* [95% CI]** |
| Intercept | **3.45 [3.44, 3.46]** | **5.78 [5.37, 6.19]** | -8.24 [-9.85, 28.36] |
| ALR | **0.005 [0.001, 0.008]** | **0.015 [0.008, 0.022]** | 0.09 [-0.11, 0.30] |
| Age | 0.003 [-0.001, 0.006] | **0.013 [0.006, 0.020]** | **0.71 [**0.11, 1.27] |
| Quadratic age | -0.001 [-0.001, 0.000] | **-0.001 [-0.003, -**  **0.0001]** | **-0.16 [**-0.30, -0.03] |
| Body mass | NA | 0.05 [-0.06, 0.17] | -3.55 [-7.88, 0.46] |
| Badge size | NA | NA | 0.33 [-1.99, 2.59] |
| **Random effects** | **σ [95% CI]** | **σ [95% CI]** | **σ** |
| Island-year | 0.008 [0.000, 0.012] | 0.000 [0.000, 0.016] | 0.739 |
| Individual | 0.057 [0.052, 0.061] | 0.011 [0.010, 0.012] | 0.413 |
| Residual variance | 0.029 [0.027, 0.031] | 0.058 [0.055, 0.062] |  |
| **Sample sizes** | **n** | **n** | **n** |
| Island-year | 170 | 170 | 112 |
| Individuals | 483 | 483 | 283 |
| Observations | 1024 | 1024 | 448 |

**Table S10:** Direct path coefficients of age (including age-at-last-reproduction ALR) and morphology on multiple paternity for females (Female MP), using only first clutches. Values in bold are supported by the model (95% CrI not overlapping zero). The results are based on the Bayesian model with sub-models described in the text.

|  | **Body mass** | **MP** |
| --- | --- | --- |
| **Direct effect** ^a^ | ***β* [95% CrI]** | ***β* [95% CrI]** |
| Intercept ^b^ | **3.45 [3.44, 3.45]** | 10.50 [-10.28, 32.08] |
| ALR | **0.01 [0.01, 0.02]** | 0.24 [0.00, 0.47] |
| Age | -0.00 [-0.01, 0.01] | 0.28 [-0.46, 1.03] |
| Quadratic age | 0.001 [-0.001, 0.002] | -0.12 [-0.28, 0.01] |
| Body mass | - | -3.49 [-9.76, 2.47] |
| Breeding synchrony | - | 0.13 [-0.24, 0.49] |
| **Compound path** |  |  |
| ALR | - | 0.20 [-0.02, 0.47] |
| Age | - | 0.28 [-0.48, 1.03] |
| Quadratic age | - | -0.12 [-0.28, -0.05] |
| **Random effects** | **σ [95% CrI]** | **σ [95% CrI]** |
| Island-year | 0.02 [0.01, 0.02] | 0.87 [0.21, 1.48] |
| Individual | 0.06 [0.06, 0.07] | 0.55 [0.03, 1.29] |
| Residual variance | 0.05 [0.05, 0.05] |  |
| **Sample sizes** | **n** | **n** |
| Island-year | 184 | 99 |
| Individual | 608 | 211 |
| Observations | 1121 | 284 |

^a^ Estimates rounded to two decimals, unless the third decimal is needed to identify positive or negative values, in

which three decimals are given ^b^ Estimates with age 1 as reference (1-year olds set to 0) for all traits

**Table S11:** Direct path coefficients of age (including age-at-last-reproduction ALR) and morphology on body size and multiple paternity for females (MP), including all clutches over the entire breeding season, using a frequentist approach with the “lme4”-package. Values in bold are supported by the model (95% CI not overlapping zero).

|  | **Body mass** | **MP** |
| --- | --- | --- |
| **Direct effect** | ***β* [95% CI]** | ***β* [95% CI]** |
| Intercept | **3.44 [3.44, 3.45]** | 2.99 [-4.97, 10.84] |
| ALR | **0.011 [0.006, 0.016]** | **0.19 [0.06, 0.34]** |
| Age | -0.001 [-0.001, 0.001] | 0.06 [-0.32, 0.43] |
| Quadratic age | 0.001[-0.001, 0.001] | -0.05 [-0.13, 0.03] |
| Body mass | NA | -1.08 [-3.36, 1.23] |
| **Random effects** | **σ [95% CrI]** | **σ [95% CrI]** |
| Island-year | 0.017 [0.011, 0.022] | 0.00 [0.000, 0.77] |
| Individual | 0.062 [0.057, 0.068] | 0.00 [0.000, 1.03] |
| Residual variance | 0.048 [0.045, 0.051] |  |
| **Sample sizes** | n | **n** |
| Island-year | 184 | 119 |
| Individual | 608 | 282 |
| Observations | 1121 | 456 |

**Table S12:** The effects of badge size and body mass on multiple paternity (MP) over the entire breeding season for males without controlling for any age component(s) using a frequentist approach with the “lme4”-package.

|  | **MP** |
| --- | --- |
| **Direct effects** | ***β* [95% CI]** |
| Intercept | 0.65 [-18.90, 19.14] |
| Badge size | 1.11 [-1.21, 3.33] |
| Body mass | -2.56 [-6.47, 1.61] |
| **Random effects** | **σ** |
| Island-year | 0.41 |
| Individual | 0.21 |
| **Sample sizes** | **n** |
| Island-year | 112 |
| Individuals | 283 |
| Observations | 448 |
